# Protection against maternal infection-associated fetal growth restriction - proof-of-concept with a microbial-derived immunomodulator OM85: safety and efficacy data

**DOI:** 10.1101/064857

**Authors:** NM Scott, JF Lauzon-Joset, AC Jones, KT Mincham, NM Troy, J Leffler, M Serralha, SL Prescott, SA Robertson, C Pasquali, A Bosco, PG Holt, DH Strickland

## Abstract

Infection-associated inflammatory stress during pregnancy is the most common cause of fetal growth restriction and/or miscarriage. Treatment strategies for protection of at-risk mothers are limited to a narrow range of vaccines, which do not cover the bulk of the common pathogens most frequently encountered. Employing mouse models, we demonstrate that oral treatment during pregnancy with a microbial-derived immunomodulator (OM85^TM^), currently used clinically for attenuation of infection-associated airway inflammatory symptoms in infants-adults, markedly reduces risk for fetal loss/growth restriction resulting from maternal challenge with bacterial LPS or influenza. Focusing on LPS exposure, we demonstrate that the key molecular indices of maternal inflammatory stress, notably high levels of RANTES, MIP-1a, CCL2, IL-8 and G-CSF in gestational tissues/serum, are abrogated by OM85 pretreatment. Systems-level analyses conducted in parallel employing RNASeq revealed that OM85 pretreatment selectively tunes LPS-induced activation in maternal gestational tissues for attenuated expression of TNF-, IL1-, and IFNg-driven that drive production of these pro-inflammatory cytokines, without constraining Type1-IFN-associated networks central to first-line anti-microbial defense. This study suggests that broad-spectrum protection-of-pregnancy against infection-associated inflammatory stress, without compromising capacity for efficient pathogen eradication, represents an achievable therapeutic goal.

**Disclosure:** This study was funded principally by Nation Health and Medical Research Council (NHMRC) of Australia with supplementary support provided by OM Pharma (Geneva, Switzerland).

CP is an employee of OM Pharma (Vifor Pharma). The other authors declare that they have no conflict of interest.

## Introduction

Systemic inflammatory processes triggered by infections during pregnancy can have adverse short term effects on maternal well-being (1, 2), and can additionally negatively impact on ensuing fetal growth and survival via disturbance of immune homeostatic processes in gestational tissues (3, 4). Moreover, intrauterine growth restriction following maternal infection is also associated with increased disease risk postnatally in surviving fetuses. A prominent recent illustration is Zika virus infection, but this represents an extreme example of a much more ubiquitous problem involving multiple (including common) pathogens, and the broader principle that fetal growth restriction regardless of cause, constitutes a risk factor for development of a range of chronic diseases in later life is now widely recognised (5, 6). With respect to risk associated specifically with infections, maternal immunization represents the only protective treatment option currently available (7), but the breadth of potential coverage against the wide range of common pathogens potentially encountered during pregnancy is severely limited by vaccine availability.

To address this latter limitation, we have turned to an alternative concept, notably the potential use of immunomodulator(s) to boost the capacity of the maternal immune system to efficiently clear infections, a capacity which is known to be compromised during pregnancy (1, 2). For this purpose, we have utilized we have turned to an orally delivered microbial-derived immunomodulatory agent OM85. This agent comprises a standardized lyophylized extract of a mixture of bacterial pathogens containing multiple TLR ligands (8, 9), and has a well-established clinical safety profile spanning a period dating back to the 1980s, including in infants as young as 6 months (10-12). In independent randomized clinical trials OM85 has previously been shown to attenuate infection-associated inflammatory symptoms in infants (13, 14) and adults (15) with predisposition to recurrent viral and/or bacterial infections. We hypothesized that prophylactic treatment during pregnancy could potentially provide broad-spectrum protection against the effects of microbial pathogens in general on pregnancy outcomes, and in the present study we have sought proof-of-concept and associated mode-of-action data in a mouse model.

## Results

### Protection against exaggerated responses to influenza infection during pregnancy

We employed a model utilizing the Influenza A/H1N1/PR8 (PR8) murine strain (16) (Supplementary Fig. 1) to investigate the effects of OM85 pretreatment on the ensuing effects of live Influenza A virus (IAV) infection (on gd 9.5) during pregnancy. In humans, pregnancy has been associated with a predisposition to IAV infection and heightened disease severity (2, 17, 18). A comparable pattern was clearly evident in infected pregnant mice 8 days after infection with PR8 with increased levels of viral replication in lung tissue (Supplementary Fig. 2a) which was associated with higher clinical distress scores (Supplementary Fig. 2b) and proportionately higher weight loss (Supplementary Fig. 2c) over the disease course relative to comparably infected non pregnant mice.

To evaluate the capacity of OM85 to protect against these effects in pregnant mice, a series of animals were pretreated with OM85 or placebo for 8 consecutive days from vaginal plug detection (gd0.5) until the day preceding infection (gd8.5). Maternal clinical data were collected daily until sacrifice at outcome day 17.5, and as illustrated in Figure 1a, OM85 pretreatment reduced clinical stress scores from day 14 onwards, but did not significantly modify infection-associated weight loss (Fig. 1b). An additional series of assessments were performed at outcome day 17.5, including cellular profiling of bronchoalveolar (BAL) washouts, PCR quantitation of viral copy numbers in lung tissue homogenates, and fetal /placental weights measured. These data together with outcome day maternal weight and clinical scores, were integrated via Principal Component Analysis (19) as summarized in Figure 1c and Supplementary Table 1, which identified two clusters within the OM treated group. One of these (cluster A) clearly demonstrated attenuated severity of infection related outcomes (maternal and fetal) with a response phenotype that falls between non-infected controls and the PR8-infected/untreated animals. The second cluster (B) did not demonstrate any overall effect of OM85 pretreatment on the response to IAV infection and data remain overlapped with the PR8-infected/non-treated group in the PCA. This heterogeneity of OM85 responder phenotypes suggested that the treatment regimen (which comprised pre-infection dosing only) may have been suboptimal. Extending OM85 treatment to include the period of infection itself markedly reduced the frequency of non-responders, and resultant clinical stress scores and maternal weight trajectories over the time course now corresponded to those of OM85 responder group A (Supplementary Fig. 3). Focusing specifically on clinical parameters on outcome day 17.5, it is evident that daily treatment significantly reduced infection-mediated clinical stress (Fig. 1d), fetal weight loss (Fig. 1e) and viral titers in lung tissue (Fig. 1f).

**Figure 1:**
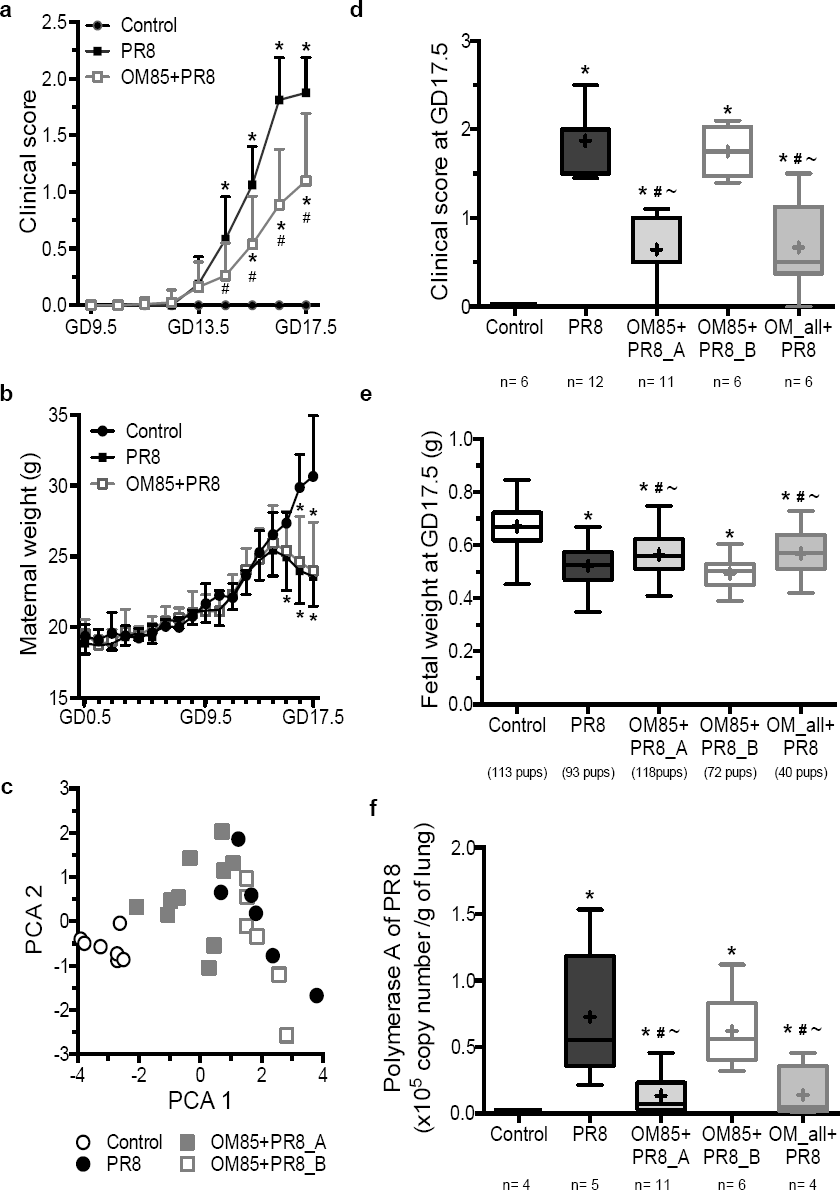
OM85 reduces the severity of Influenza infection in pregnant mice. a) Clinical score and (b) maternal weight of control non-pregnant mice versus timed mated mice infected with PR8 with or without OM85 pretreatment from gd0.5 to gd8.5. Data from mice treated with OM85 alone was not different from the Control group. (Control, n = 6; PR8, n = 12; OM85+PR8, n = 20; collected from >5 independent experiments). Data are displayed as mean ± SD. Statistical analysis was done by two-way ANOVA with Bonferroni multiple-comparison test; *: p<0.05 vs Control, #: p<0.05 vs PR8. (c) Principal component analysis (PCA) using the parameters described in Supplementary Table 1. (Control, n = 6; PR8, n = 6; OM85+PR8_A, n = 10; OM85+PR8_B, n = 6; collected from >5 independent experiments). (d,e,f) Influenza infection of mice pretreated with OM85 from gd0.5 until gd8.5 (OM85+PR8) or during the whole pregnancy (OM_all+PR8, n = 6). At gd17.5, clinical score (d), fetal weights (e) and lung viral load (measured in lung homogenate by qPCR) were assessed. The viral load of the controls was below the detection limit of the assay. Statistical analysis was done by one-way ANOVA with Bonferroni’s multiple-comparison test; *: p<0.05 vs Control, #: p<0.05 vs PR8, ~: p<0.05 vs OM85+PR8_B.

### Protection of pregnancy against the toxic effects of bacterial LPS challenge

We next employed a rigorously validated LPS exposure model (20, 21) to mimic bacterial infection during pregnancy and evaluated the potential of OM85, administered daily from gd9.5 until just prior to LPS administration on gd16.5 (Supplementary Fig.4), to attenuate LPS induced fetal resorption and/or growth restriction over the ensuing 24 hours. The dose of LPS used in this study (10ug) was selected to result in approximately 50% fetal loss with significant weight reduction (mean 13%) in surviving pups compared to those from control pregnant mice (Supplementary Fig. 5a, b and Fig. 2a, b), thus allowing for sufficient viable fetal tissues for further analyses and the potential to identify significant differences in fetal weights. Pretreatment of pregnant mice with OM85 prior to LPS administration resulted in significantly improved fetal survival rates (Fig. 2a) and prevention of fetal weight loss (Fig. 2b) compared to LPS injected control pregnant mice. We did not observe any significant changes in placenta weight across any of the groups of mice regardless of OM85 treatment or LPS injection (Fig. 2c). However, as expected due to the differences in fetal weight, the fetal:placenta weight ratio was significantly different between treatment groups (Fig. 2d, Supplementary Fig. 5d). The analyses in Fig. 2b,c,d utilized data only from pregnant mice with a litter size of 7 across all groups of mice, in this way eliminating litter size as a confounder in analyses relating to fetal weight loss; however importantly, comparable findings were obtained across the collective study when all litters were included (Supplementary Fig. 5c, d).

**Figure 2:**
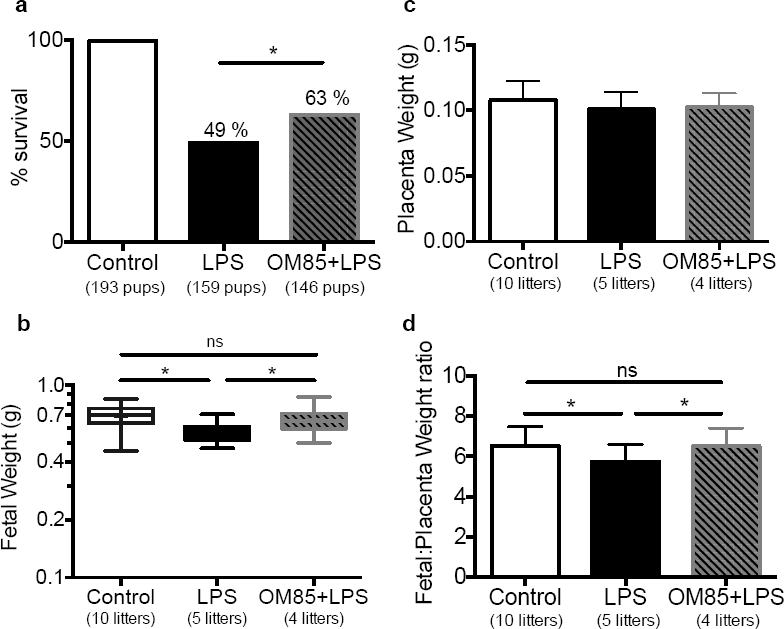
OM85 treatment prior to maternal LPS exposure protects against adverse pregnancy outcomes. (a) Percent survival of offspring from all litters size (with n>25 litters per group, total pup number per group shown in parentheses). Statistical analysis was done by Chi-squared test; *: p<0.05. (b,c,d) Time-mated mice were pretreated of not with OM85 and injected with LPS at gd16.5. At gd17.5, (b) fetal weight, (c) placenta weight and (d) fetal:placenta weight ratio were measured. Data shown are mean ± SD for litter size of 7 pups (derived from the same animals as per (a)). Statistical analysis was done by oneway ANOVA with Bonferroni multiple-comparison test; *:p<0.05, ns=not significantly different.

### Modulation of infection-associated immune cell trafficking in gestational tissues during pregnancy

Using the LPS model, we next proceeded to studies on the effects of OM85 treatment during pregnancy on immune cell populations known to play important roles in the maintenance of immunological homeostasis in gestational tissues (22-25), using multi-parameter flow cytometry (detailed in methods, panels shown in Supplementary Table 2 and gating strategy described in Supplementary Fig. 6 and 7). The gestational tissues analyzed are referred to in the text as placenta, uterus, decidua (see Methods and Supplementary Figure 4 for a more detailed description of precise tissue collection methodology) and draining lymph nodes (PALN). Total CD45^+^ cell yields were not significantly influenced by LPS exposure (data not shown). Amongst CD45+ cells, the functions of several subpopulations are known to be critically important to successful pregnancy (23) and LPS induced occasional proportional changes for total myeloid, T-cell, B-cell, and macrophage populations in the tissues examined, but these were not modified by OM85 pretreatment (Supplementary Fig. 8). However, as illustrated in Figure 3, the profile of the cellular response to LPS exposure in animals pretreated with OM85 was notably different for four rare immunomodulatory cell populations: T-regulatory (Treg), plasmacytoid DCs (pDC), conventional DCs (cDC) and myeloid derived suppressor (MDSC) cells. In our model, LPS drives marked accumulation of MDSCs in uterus/placenta (Fig. 3a,b) and Tregs in uterus/decidua (Fig. 3a,c). Induction of this inflammatory cellular response was attenuated in OM85 pretreated pregnant mice; dysregulation of MDSC at the fetomaternal interface has been associated with poor pregnancy outcomes (26). In contrast, LPS exposure depleted resident uterine/placental cDC populations (Fig. 3a,b), likely by stimulating their migration to draining PALN (Fig. 3d), and this was also attenuated by OM85. Moreover, the LPS-induced upregulation of activation markers CD40 and CD86 on cDC in gestational tissues was conserved and/or enhanced by OM85 (Supplementary Fig. 9). It is pertinent to note in this regard that entrapment of functional cDC in gestational tissues has been linked to regulatory processes that contribute to successful pregnancy outcomes (27). It is additionally evident that recruitment of pDC into uterus/decidua by LPS is significantly enhanced by OM85 pretreatment (further discussion below).

**Figure 3:**
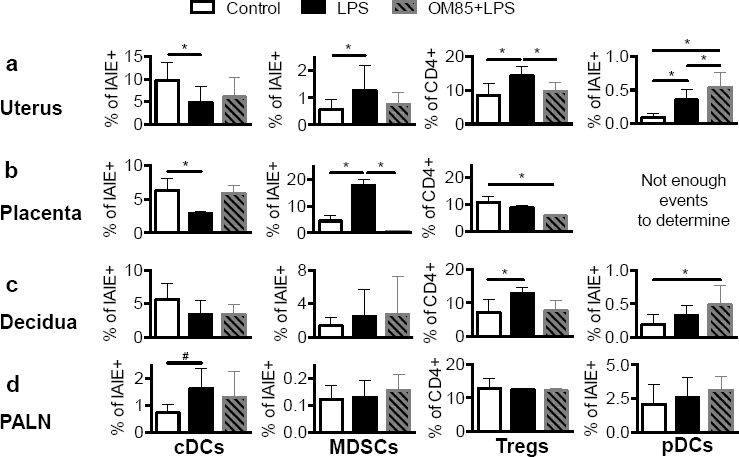
OM85 pretreatment modulates the inflammatory immune cell response to maternal LPS exposure. Single cell suspensions prepared from (a) uterus, (b) placenta, (c) decidua, and (d) para-aortic lymph node (PALN) tissues collected from control pregnant versus LPS exposed pregnant mice with and without OM85 pretreatment, were analyzed via flow cytometry for cDCs (CD45^+^F480^−^I-A/I-E^+^Ly6G/C^−^B220^−^CD11c^+^), MDSCs (CD45^+^F480^−^I-A/I-E^+^Ly6G/C^+^B220^−^CD11c^−^), Tregs (CD45^+^CD3^+^CD4^+^CD8^−^CD25^+^FoxP3^+^) and pDCs (CD45^+^F480^−^I-A/I-E^+^Ly6G/C^+^B220^+^CD11b^−^CD11c^lo^). Data displayed as mean ± SD, n = 7 per group collected from >5 independent experiments. Statistical analysis was done by one-way ANOVA with Bonferroni multiple-comparison test (*: p<0.05) or by unpaired t-test (#: p<0.05).

### Attenuation of infection-associated inflammatory mediator production in gestational tissues

As measures of maternal inflammatory stress, we next determined levels of a broad array of cytokines in gestational tissues as above and maternal serum at the time of sacrifice on gd17.5 (Supplementary Fig. 10,11,12). The largest and most consistent changes induced by LPS involved RANTES, MIP-1a, KC (IL-8), CCL2 (MCP-1) and G-CSF production. Of note, the production of these pro-inflammatory mediators in response to LPS was considerably reduced in pregnant mice that were pretreated with OM85 (Fig. 4a,b,c for uterus, placenta and maternal serum respectively). The observation that some but not all pro-inflammatory cytokines were attenuated at the protein level suggested a degree of selectivity in the action of OM85 pretreatment in orchestrating the direction of the immune response to LPS. To explore this finding in more detail at the systems-level, we employed RNASeq profiling of maternal tissues.

**Figure 4:**
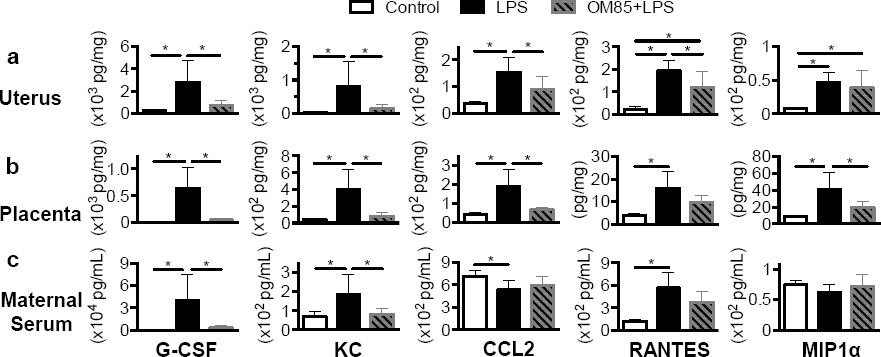
OM85 pretreatment prior to maternal LPS exposure attenuates the inflammatory cytokine response. Protein extracted from (a) uterus (b) placenta and (c) maternal serum collected at gd17.5 from control pregnant versus LPS exposed pregnant mice with and without OM85 pretreatment was used to determine tissue cytokine profiles. Shown are the levels of G-CSF, KC, MCP-1, RANTES, and MIP1? as assessed by multiplex cytokine analysis. Data displayed as mean ± SD, n = 5-6 per group collected from >3 independent experiments. Statistical analysis was done by one-way ANOVA with Bonferroni multiple-comparison test; *: p<0.05.

### Systems level analyses targeting mechanism-of-action of OM85

To decipher the molecular mechanisms that underpin OM85 mediated reprogramming of the LPS response, we identified differentially expressed genes and networks (28, 29) from RNASeq profiles of the uterus (using portions of same samples analyzed for cytokine levels and cellular profiling) and decidua (obtained from independent animals from each group). These data were further interrogated with upstream regulator analysis (30) to identify the molecular drivers of the responses (detailed in methods). LPS challenge induced a strong perturbation of the gene expression program in decidua and uterus compared to control (non LPS-exposed) pregnant mice. Specifically, in the order of 3000 differentially expressed genes (DEG) were identified in each tissue (FDR <0.05, decidua: Fig. 5a and Supplementary Table 3 and uterus: Supplementary Fig. 13a and Supplementary Table 4), and subsequent pathways analyses employing InnateDB clearly identified a series of LPS-sensitive innate and adaptive immunoinflammatory pathways in both tissues (decidua: p values 3 × 10^−1^ 1.2×10^−21^, Supplementary Table 5, uterus: p values 2.5×10^−7^ - 6×10^−19^, Supplementary Table 6).

**Figure 5:**
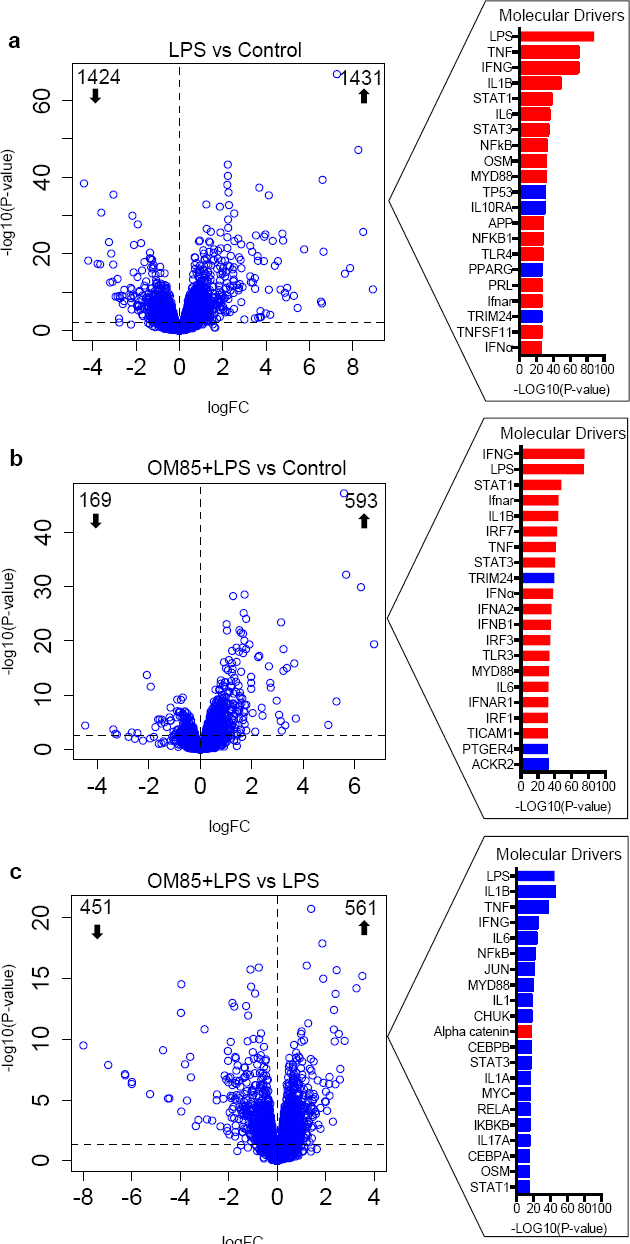
OM85 pretreatment prior to maternal LPS exposure attenuates inflammatory gene programs in decidua. Differentially expressed genes were identified using edgeR in decidua from (a) LPS challenged versus control pregnant mice; (b) OM85 pretreated LPS challenged versus control pregnant mice; (c) OM85 pretreated LPS challenged versus LPS challenged pregnant mice. Data are mean, n=5 per group collected from >3 independent experiments. The dashed horizontal lines indicate an FDR<0.05. Right panels: Molecular drivers of the differential expression patterns were identified using Upstream Regulator Analysis. Driver genes shown in red were activated and those shown in blue were inhibited (all drivers had absolute activation Z-scores > 2.0).

Parallel analyses comparing gene expression in LPS-challenged OM85-pretreated pregnant mice to controls revealed a marked decrease in the number of genes induced by LPS, with 762 DEG identified in decidua (FDR<0.05, Fig. 5b, Supplementary Table 7) and 709 DEG in uterus (Supplementary Fig. 14b, Supplementary Table 8) respectively, but the overall spectrum of underlying LPS-sensitive immunoinflammatory pathways (decidua: p values 10^−7^ − 1.2×10^−28^, Supplementary Table 9; uterus: p values 10^−10^ − 10^−25^, Supplementary Table 10) was qualitatively similar to that seen above in tissues from non treated animals. However, a direct comparison between LPS challenged mice with versus without OM85 pretreatment highlighted significant treatment-associated attenuation of multiple inflammation-associated genes in these tissues (exemplary genes in Supplementary Fig. 14; global responses in Fig. 5c, Supplementary Fig. 13c and Supplementary Tables 11/12).

Next, we employed upstream regulator analysis to identify the putative molecular drivers of the observed differential gene expression patterns. These analyses confirmed LPS itself as the most significant driver of the LPS response (Fig. 5a), demonstrating the overall plausibility of this model with the published LPS literature. At the top of the rank order of subsequent drivers of the LPS response were TNF, IFNG, IL1B and IL6, which respectively accounted for 422, 367, 250 and 196 of the differentially expressed genes in the decidua. Similarly in the uterus we identified IFNG, TNF, IL1B, and IL6 as the most significant drivers of the LPS response, which respectively accounted for 390, 428, 264, 206 of the differentially expressed genes (Supplementary Fig. 13a). Consistent with reduced inflammatory responses to LPS in mice pretreated with OM85, the number of genes for the same drivers was dramatically reduced in the decidua (TNF: 152, IFNG: 179, IL1B: 114 and IL6: 89; Fig. 5b) and in the uterus (IFNG: 192, TNF: 154, IL1B: 110, IL6: 86; Supplementary Fig. 13b). However, it is noteworthy that in the decidua and the uterus of OM85 pretreated mice, genes from the type I interferon pathway were enriched amongst the top of the rank order of molecular drivers (IFNAR1, IRF7, IFNA, IFNB; Fig. 5b and Supplementary Fig. 13b), whereas proinflammatory drivers (TNF, IL-1B, IL-6) in general had lower ranks. Lastly, we identified the molecular drivers of the differential response to LPS challenge in mice with or without OM85 pretreatment. The data showed that the expression of multiple proinflammatory pathways (IL1B, TNF, IFNG, and IL6; Fig. 5c) was decreased in the decidua of OM85 pretreated mice after LPS challenge in comparison to LPS treated control pregnant mice. We observed similar patterns of attenuated inflammatory gene expression programs in the uterus (IFNG, TNF, IL1B and IL6, Supplementary Fig. 13c).

We employed coexpression network analysis to provide a holistic view of the gene expression program in decidua and uterus, focusing on the most variables genes. In the decidua, the resulting network contained 4271 genes organized into 17 coexpression modules (Fig. 6a). LPS challenge modulated the expression of seven modules (modules A, D, E, F, H, J and Q; median FDR < 0.05, Fig. 6b). In contrast, if the mice were pretreated with OM85, only two modules were perturbed by LPS challenge (modules A and F; median FDR < 0.01, Fig. 6c). In the uterus, the LPS-induced coexpression network comprised 4185 genes structured into 11 coexpression modules (Fig. 6d). LPS challenge perturbed the expression of four modules (B, I, J and K; FDR<0.05, Fig. 6e), but only one of these modules was modulated by LPS in mice pretreated with OM85 (module B; median FDR< 0.01, Fig. 6f). Lastly, we characterized the upstream regulators that coordinated the effect of each differentially expressed module in both tissues (Fig. 7 and Supplementary Fig. 15). In decidua, the LPS response was largely driven by proinflammatory pathways (e.g. OSM, TNF, IL-1B, CEBPB, NF-kB, MyD88, module E, module J, module Q; Fig. 7a). OM85 pretreatment silenced the proinflammatory upstream regulators while conserving the modules driven by IRF7 (module A) and IFNG (module F) (Fig. 7b). Similar effects were seen in the uterus, upstream regulator analysis suggested that module B was driven by IFNG and IRF7 (Supplementary Fig. 15a,b). Again, OM85 pretreatment silenced modules associated with proinflammatory pathways (e.g. TNF, IL-1B, IL-6, NF-kB, module I, module K; Supplementary Fig. 15a), but not Type I IFN response pathways (Fig. S14). These data show that OM85 pretreatment preferentially attenuates proinflammatory gene networks, but preserves interferon-mediated networks.

**Figure 6:**
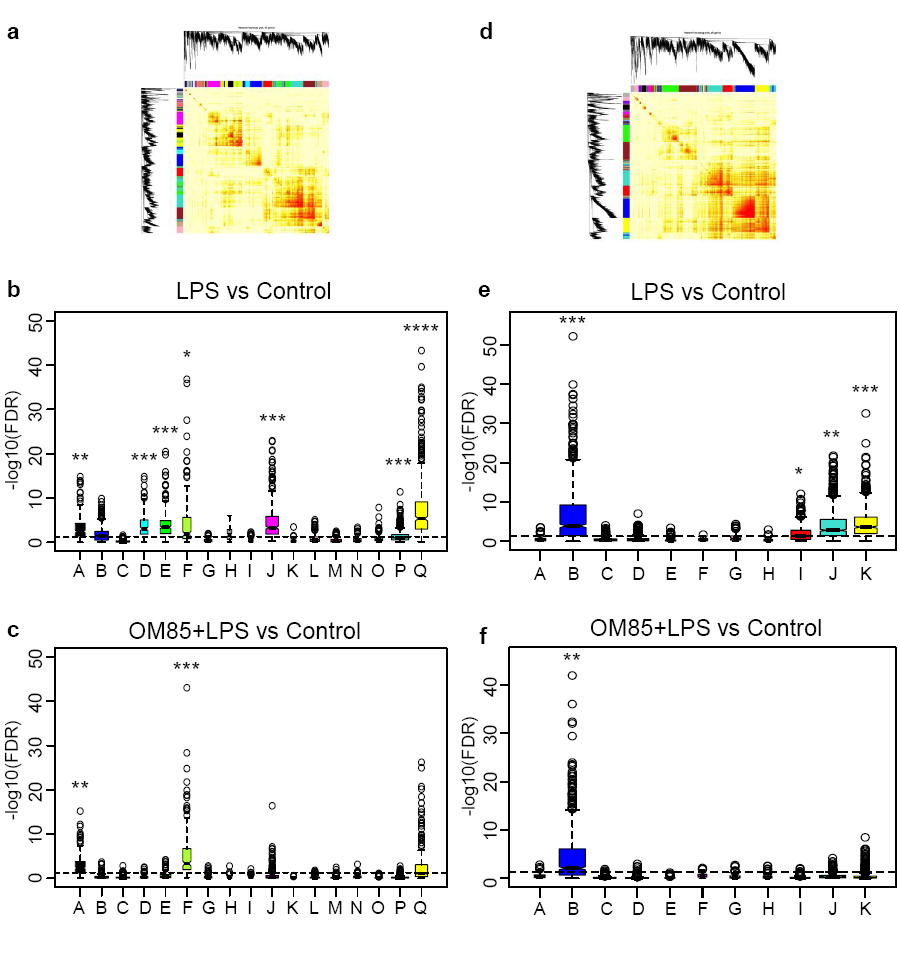
OM85 pretreatment modulates expression of inflammatory gene coexpression networks in decidua and uterus following maternal LPS exposure. Network analysis (WGCNA) was performed on gene expression patterns focusing on the most variable genes. Coexpression network topology in decidua (a) and uterus (d). Red block-like structures indicate modules of coexpressed genes. The overall expression of the modules was compared in LPS challenged versus control pregnant mice in decidua (b) and uterus (e); and in OM85 pretreated LPS challenged versus control pregnant mice in decidua (c) and uterus (f). The p values were derived from an edgeR analysis and the dashed horizontal lines indicate an FDR<0.05. ****: median FDR <0.0001; ***: median FDR <0.001; **: median FDR <0.01; *: median FDR <0.05.

**Figure 7:**
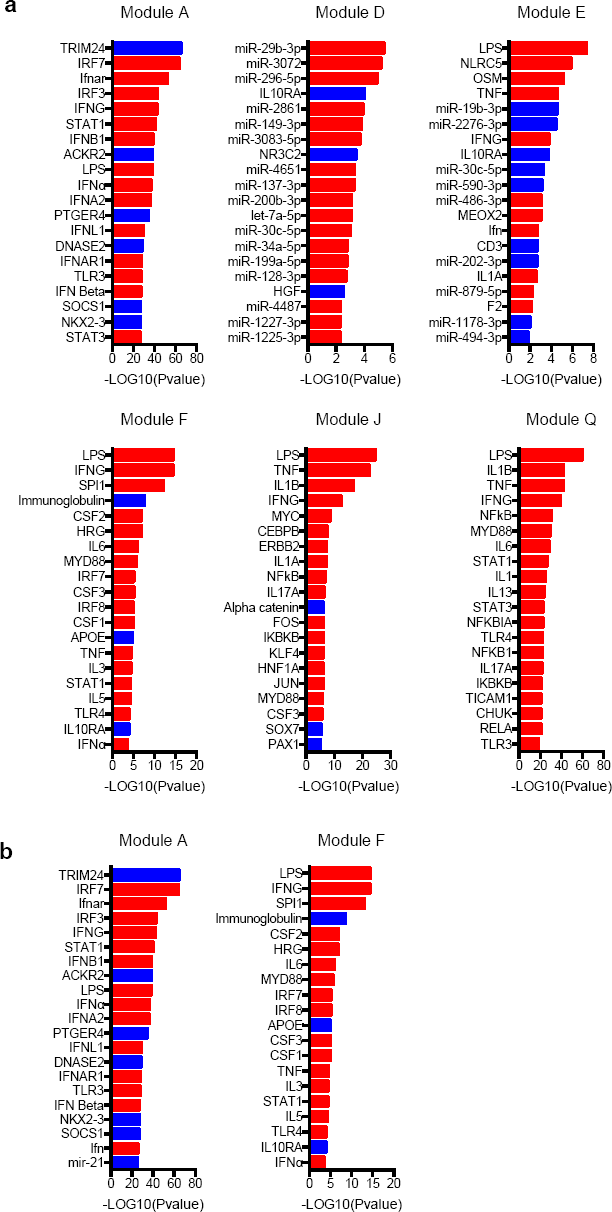
Molecular drivers of gene coexpression modules in decidua from LPS challenged mice with or without OM85 pretreatment versus control pregnant mice. Molecular drivers of gene coexpression networks in decidua of (a) LPS challenged versus control pregnant mice; (b) OM85 pretreated LPS challenged versus control pregnant mice. Molecular drivers highlighted in red denote activation and blue indicates inhibition.

### Safety profile of OM85 during normal pregnancy

In conjunction with the functional studies detailed above relating to OM85 use as an anti-inflammatory agent, we additionally evaluated the effects of prophylactic treatment with OM85 during gestation on normal pregnancy outcomes in the absence of infectious challenge. RNASeq analyses demonstrated that groups of pregnant mice exposed only to OM85 treatment for 8 days from gd9.5 (as per LPS model) induced a very limited perturbation of the gene expression program with only 15 and 1 differentially expressed genes (DEG) seen in the decidua and uterus respectively (Supplementary Fig. 16). In terms of cellular composition within gestational tissues, the proportions of the immune cell populations including NK cells, cDC, pDC, B cells, T regulatory cells, MDSC and other myeloid populations were unchanged by OM85 treatment (Supplementary Fig. 17), with the exception of small but significant increases in frequency of macrophages and T-cells in the placenta only (Supplementary Fig. 17b). OM85 treatment did not skew the cytokine milieu across the range of Th1, Th2, proinflammatory and regulatory cytokines measured in the maternal serum, uterus and placental tissue (Supplementary Fig. 18). On average OM85 decreased cytokines in the maternal serum by 20%, the uterus tissue by 5%, and increased cytokines the placental tissue by an average of 36% (Supplementary Fig. 18).

Groups of control pregnant mice versus those treated for 8 days from gd 9.5 with OM85 (as per the LPS model) had similar weight gain curves (Fig. 8a) and litter sizes (Fig. 8b). Additionally, there were no significant differences in the mean pup birth weight or growth trajectory (Fig. 8c), fetal or placenta weight or fetal:placenta weight ratio at gd17.5 (Fig. 8d,e,f respectively). Further, we did not find any differences at gd17.5 in number of viable implantations per pregnant mouse, gestational time to parturition, number of resorptions per pregnant mouse, or fetal weights (Supplementary Fig. 19a,b,c,d respectively) between groups of control versus treated pregnant mice. In separate experiments we also determined the safety of OM85 given over the first 8 days from detection of plug as per the Influenza model on the clinical outcomes of pregnancy and as above found no changes (data not shown). Collectively, across maternal and fetal clinical parameters, immune cell and molecular profiling, our data provides supporting evidence that OM85 can be safely used during gestation.

**Figure 8:**
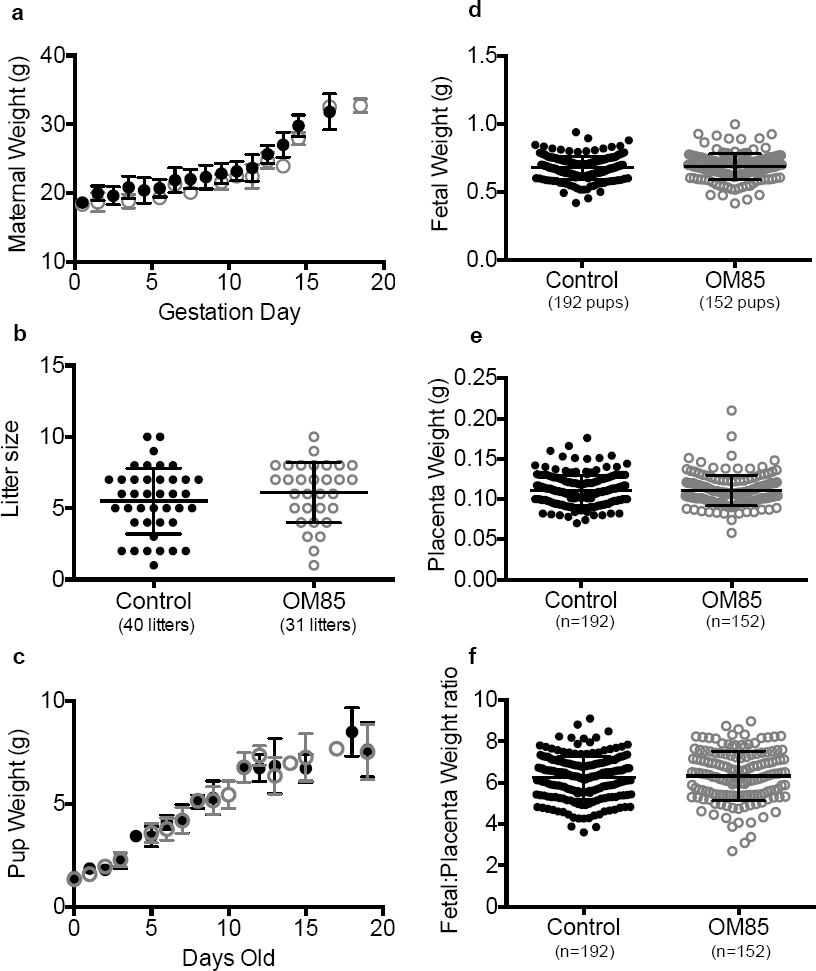
OM85 treatment did not alter normal pregnancy outcomes. Timed mated female mice treated with OM85 or naïve/vehicle treated controls were followed through pregnancy. (a) Maternal weight trajectory during pregnancy (control n=26 pregnancies, OM85 n=13 pregnancies), (b) litter size, and (c) pup weight trajectory from birth until weaning (control n=11 litters, OM85 n=19 litters) were measured. Some mice were autopsied at gd 17.5 and (d) fetal weight, (e) placenta weight, and (f) Fetal:placental weight ratio were assessed. Data displayed as mean ± SD, number of pups or litters shown in parentheses; no significant differences were found by unpaired t test.

## Discussion

Immune function(s) in gestational tissues require fine control to balance the conflicting needs of suppression of maternal responses against fetal allograft antigens, while enabling effective defense against microbial pathogens. An additional imperative is that maternal expression of antimicrobial immunity, particularly elements of these responses that have potential to spill over into the systemic compartment, must be tightly regulated to limit the risk of “bystander” inflammatory damage to the highly vascularized tissues at the fetomaternal interface. However this balance is frequently not achieved, and indeed heightening of infection susceptibility and accompanying symptomatology is a recognized feature of the normal pregnant state (2, 24, 31). This infection-susceptible phenotype is additionally a common feature of infancy, a hallmark of which is the severe lower respiratory tract infections which peak in frequency within the first year of life (32, 33). Recent clinical studies suggest that prophylactic treatment of infants with the microbial-derived immunomodulatory agent OM85 can significantly attenuate the intensity and duration of their respiratory infection-associated symptoms (10-14). We posited that by extension, this same agent may offer similar protection during pregnancy, and if so this may also mitigate the known downstream effects of maternal infection on fetal growth restriction and/or loss.

Employing a model of live IAV infection with an inoculum which permitted 100% survival of pups over the ensuing 8 days, pretreatment with OM85 resulted in a dose-dependent enhancement of IAV clearance in infected pregnant mice, significantly reduced maternal clinical stress scores across the infection time course, and attenuated the infection-associated inhibition of fetal growth rates, as shown in Figure 1/Supplementary Figure 3.

OM85 pretreatment also provides significant protection, as illustrated in Figure 2, against the effects of maternal bacterial LPS exposure on subsequent acute fetal loss and/or growth restriction. Mechanistic studies focusing on the LPS model to characterize the nature of the changes induced by OM85 pretreatment in association with protection against the toxic effects of LPS revealed the capacity of OM85 to fine tune the immune response. Systems-level analyses of RNASeq profiles from maternal gestational tissues provided a global view of the underlying gene networks involved. This demonstrated that LPS exposure of control pregnant mice perturbed around 3,000 genes in the decidua and uterus, which was markedly reduced in mice pretreated with OM85 to around 700 genes. Upstream regulator analysis suggested that the LPS response was mainly driven by TNF, IL1B, IL6, and IFNs, and OM85 pretreatment selectively attenuated the activation of proinflammatory pathways (TNF, IL1B, IL6) whilst the interferon response pathway (IRF7, IFNG) (34-36) was preserved. It is noteworthy that TNF, IL1 and IL6 are prominent amongst the list of inflammatory mediators that are recognized contributors to pregnancy loss/complications (37, 38) and accordingly selective suppression of their production while concomitantly conserving type 1 IFN pathways that are central to anti-microbial defense, provides a plausible explanation for the pregnancy sparing effects of OM85 pretreatment in the face of microbial challenge.

The precise molecular mechanism(s) through which OM85 regulates the balance between these inflammatory and microbial defense pathways following LPS binding remains to be determined. A precedent for differential regulation of these two pathways can be found in recent literature demonstrating that LPS triggering can sequentially activate two distinct signalling pathways, via plasma membrane-localized and endosomal TLR4/LPS complexes (35, 39). The former induces TIRAP-MyD88 signalling while the latter induces TRAM-TRIF signalling, resulting respectively in production of pro-inflammatory cytokines and type 1 IFNs. On this basis it is feasible that one consequence of OM85 treatment may be to alter the balance between cell surface versus intracellular TLR4/LPS signalling in target cells, and this possibility is amenable to testing.

The mode of action of OM85 parallels the concepts of trained immunity, a term which describes the augmentation of innate immune function following a stimulus not specific to the original stimulus (32, 40, 41). Further examples of the latter include protection against unrelated infectious diseases through vaccination and the seminal studies that have described striking protective effects of environmental microbial exposure via inhalation and dietary intake during pregnancy through living/working in a traditional European farming environment, on (*inter alia*) allergy and asthma development in offspring (42, 43).

LPS exposure of pregnant mice induces high levels of inflammatory cytokine/chemokine production (Fig. 4) accompanied by recruitment of anti-inflammatory immune cell populations (Tregs and MDSCs; Fig. 3) in 2 of the 3 gestational tissues tested. Both mediator production and immune cell recruitment were attenuated in OM85 pretreated mice, consistent with reduction of the local inflammatory burden. An additional feature of the LPS response was depletion of resident cDC in gestational tissues (Fig. 3a,b) which, in common with other inflammatory challenge models (44, 45), likely involves stimulation of their migration to draining lymph nodes in response to inflammatory triggers (Fig. 3d). This depletion was blocked in OM85 pretreated mice (Fig. 3a,b) suggesting that reducing the proinflammatory signature prevents egress of resident DC, or promotes recruitment of fresh cDC precursors to replace the LPS-responsive emigrant population, and moreover enhances their activation upon arrival (Supplementary Fig. 9a-c). In this regard, it is noteworthy that LPS exposure additionally triggered local infiltration by pDC which are important components of the innate immune response to pathogens (46, 47), and this recruitment was also markedly enhanced in OM85 pretreated mice (Fig. 3a,c), suggesting that the overall DC precursor compartment represents a major OM85 target.

With respect to the associated cellular effector mechanisms, it is pertinent to note that the cDC and pDC populations in gestational tissues identified above as prominent OM85 targets are both recognized as key contributors to regulation of pathogen specific immunity (46, 48, 49). Moreover, upregulation of cDC function has previously been identified in association with OM85 mediated stimulation of protective antibody responses against respiratory pathogens in a mouse model (50).

In conclusion, this study provides experimental proof-of-concept that oral treatment during pregnancy with the microbial derived agent OM85 provides broad spectrum protection against the systemic effects of inflammatory responses triggered by both bacterial and viral agents, in particular against responses in gestational tissues that are associated with fetal loss and/or growth restriction. These findings suggest that existing microbial derived therapeutics with immunomodulatory properties and proven safety records may provide novel therapeutic options for protection against the toxic effects of infections during pregnancy. Moreover, they point towards opportunities for development of defined molecular entities with comparable modes-of-action, for therapeutic use in this and related clinical contexts.

## Methods

### Animals

Specific pathogen free BALB/c mice were obtained from the Animal Resources Centre (Perth, WA, Australia). Female mice were used for timed mating only between 8-12 weeks of age. Male studs were used from 8 weeks of age and retired at 36 weeks of age. All mice were housed under specific pathogen free conditions at the Telethon Kids Bioresources Facility, with food and water *ad libitum* and a 12 h light/dark cycle. All experiments were approved by the Telethon Kids Animal Ethics Committee, and strictly conducted according to the NHMRC guidelines for the use of animals for scientific research.

### Time Mated Pregnancy

Male stud mice were caged individually. Up to 2 female mice were placed in a male cage overnight and the following morning females were separated from the males and checked for vaginal plugs as evidence of mating. Females were designated gestational day (gd) 0.5 on the day of vaginal plug detection and housed in groups of 5-10 until commencement of treatments. Females not pregnant after plug detection were euthanized.

### Treatment protocols

#### OM85 Treatment

OM85 was administered orally via pipette at a dose of 400mg/kg body weight in PBS per day. In the Influenza A virus infection model, time mated females were administered OM85 daily for 9 days from gd0.5 until day of PR8 infection (Supplementary Fig. 1). Control mice were left untreated, or were administered the vehicle orally on the same treatment regimens. In the LPS challenge model, pregnant female mice were administered OM85 daily from gd9.5 until LPS challenge on gd16.5, or delivery (Supplementary Fig. 4). All treatments were performed using a single batch of OM85, supplied by OM Pharma (Geneva, Switzerland).

#### Influenza infection

The mouse-adapted influenza A/PR/8/34 virus was from the American Type Tissue Culture Collection and prepared in allantoic fluid of 9-day old embryonated hens eggs. Stock virus was sub-passaged through Mardin-Darby canine kidney (MDCK) cells in Dulbecco’s modified Eagle’s medium (DMEM; Gibco, Sydney, Australia), harvested as tissue culture supernatant and viral titres determined by cytopathic effects on MDCK cells and expressed as the mean log10 tissue culture infective dose that kills 50% of the cells (TCID50) over a 5-day incubation period (16). Non-pregnant control and gd9.5 pregnant mice were inoculated intranasally (i.n.) under light inhalation isoflurane anaesthesia with a dose of 20 TCID_50_ of PR8, diluted in PBS, in a total volume of 25µl. Mice were monitored daily for weight loss and clinical score (as below) (16). At peak of disease (gd17.5), BAL and tissues were collected and fetal/placental weights were recorded (Supplementary Fig. 1).

Viral load post infection was measured in lung homogenate by real time qPCR. Lung were homogenized in PBS (10%w/v) and RNA was extracted using TRIzol (Ambion, Life Technologies, Mulgrave, VIC, Australia) and RNeasy MiniElute kit (Qiagen Gmbh, Hilden, Germany). cDNA was prepared with Quantitect Reverse Transcription Kit (Qiagen) and PR8 Polymerase A was detected using Quantifast SYBR Green PCR master mix (Qiagen) and the following primers, 5’-CGGTCCAAATTCCTGCTGA-3’ and 5’-CATTGGGTTCCTTCCATCCA-3’ (Sigma Aldrich). Copy numbers were calculated using a standard curve of known amounts of amplified cDNA.

#### LPS Challenge

Female mice at gestational day 16.5 were administered 10 µg of LPS (*Salmonella typhimurium*, Sigma Aldrich, St Louis, MO, USA) in 200 µl of PBS via intraperitoneal (i.p.) injection as previously described (21). Controls were administered 200 µl of PBS i.p. Twenty-four hours later at gestational day (gd) 17.5, an autopsy was conducted and tissues and serum collected for further analysis (Supplementary Fig. 4).

#### Animal Monitoring and Clinical Assessments

Mice were weighed daily during the acute period of infection (d0 to autopsy day). Clinical disease scores were also assessed according to the following criteria:

Score 0 - Normal appearance, healthy and active;

Score 1 - Barely ruffled fur, mildly/intermittent hunched appearance and otherwise healthy;

Score 2 – Moderately ruffled fur, elevated respiratory rate, hunched appearance with a crab-like gait, intermittent stillness and reduction of curious behavior;

Score 3 - Ruffled fur, labored breathing, hunched appearance with a crab-like gait and unresponsive to stimuli.

### Tissue Dissection

At gestational day 17.5 tissues were collected only from implantation sites from overtly normal fetuses, dead fetuses were excluded. Following euthanasia of the pregnant mouse, placentas were peeled/blunt dissected from the maternal tissue. Then, using small scissors, a portion of mesometrial uterus approximately 3mm^2^ was dissected, which was full thickness, and therefore included the decidua, but also the residual lymphoid aggregate of pregnancy and some myometrium, this we have termed ‘decidua’. The remaining anti-mesometrial uterus was also collected, which was also full thickness, and but was primarily myometrium, this we have termed ‘uterus’. The terms ‘decidua’ and ‘uterus’ have been used throughout for simplicity for the general reader. Tissue harvesting was performed consistently across all experimental groups.

### Cell Preparations

Single cell suspensions of uterus, decidua, placenta and para-aortic lymph nodes (PALN) were prepared by enzymatic digestion using methodology as previously described (16). Briefly, following dissection the tissues were sliced into small pieces, resuspended in GKN (11 mM D-glucose, 5.5 mM KCl, 137 mM NaCl, 25mM Na2HPO4) +10%FCS (Serana, Bunbury, WA, Australia) containing collagenase IV (Worthington Biochemical Corporation, Lakewood, NJ, USA) and DNase (Sigma Aldrich) and incubated at 37°C with gentle agitation as follows; Uterus: 1.5mg/ml collagenase IV and 0.2mg/ml DNase for 60 min; Decidua and Placenta: 0.75mg/ml of collagenase and 0.1mg/ml of DNase for 60 min; and PALN: 0.75mg/ml of collagenase and 0.1mg/ml of DNase for 30 min. Following digestion, tissues were finally redispersed via pipetting and debris removed by passing suspensions through cotton wool columns. Cells were pelleted and resuspended in PBS with 0.1% Bovine Serum Albumin (Bovagen Biologicals, Victoria, Australia), prepared for total cell counts or other assays as below. For the placenta, the digested preparation was resuspended in GKN+5%FCS and further enriched for leukocytes via Histopaque (Sigma Aldrich) gradient enrichment as per the manufacturer instructions.

Broncho-alveolar lavage fluid (BALF) was harvested by slowly infusing and withdrawing 1 ml of PBS containing 1mg/ml BSA from the lungs three times. The cells were pelleted and prepared for total cell counts and differential cell counts as previously described (16). Briefly, the percentage of each cell type as identified by Diff Quik stain (macrophage, neutrophil, eosinophil, lymphocyte) was calculated as a proportion of at least 300 counted cells, and this figure used to derive total numbers of each subset based on the total BALF cell count.

### Flow Cytometry

Immunostaining of viable single cells was conducted as previously described (16). Two panels of monoclonal antibodies (Supplementary Table 2) were developed to identify leukocytes of myeloid (including CD45, I-A/I-E, F480, Ly6G/C, B220, CD11c, CD11b, CD103, CD8, CD40, CD86, and NKp46 (BD Pharmingen, San Jose, CA, USA or eBiosciences, San Diego, CA, USA) and lymphoid (using CD45, CD3, CD4, CD8, CD19, CD25, CD69, Ki67 and FoxP3 as per Supplementary Table 2) (BD Pharmingen or Biolegend, San Diego, CA, USA) lineages, and subsets therein. Intracellular staining for FoxP3 was conducted using the eBiosciences FoxP3 intracellular staining buffer set. Data was collected on a 4-laser LSRFortessa flow cytometer (BD Biosciences, San Jose, CA, USA), and analyzed using FlowJo software (Version 10.0.7, Tree Star Inc, Sanford, CA, USA). The gating strategies are illustrated in Supplementary Fig. 7 (26, 51-54).

### viSNE methods

Placenta sample FCS files, with software compensation applied, were uploaded to Cytobank software (Cytobank Inc. Mountain View CA, USA), and analyzed using established methods (55, 56). The software transformed the data to arcsinh scales with cofactors ranging from 20-2500. Equal cell numbers were analyzed from each FCS file. The antibodies listed as per Supplementary Table 2 for dendritic cell and subsets identification were used to create viSNE maps using a total of 22570 cells per sample (Supplementary Fig. 8). The antibodies for Tcell and Treg identification were used to create viSNE maps using a total of 17829 cells per sample (Supplementary Fig. 8).

### Multiplex cytokine analysis on Maternal Serum

Maternal blood was collected by cardiac puncture during autopsy on gestational day 17.5. Serum was assayed for cytokines using a Bio-Plex Pro mouse cytokine 23plex kit (BioRad Laboratories Inc, Hercules, CA, USA), following the manufacturer instructions. IL-1α, IL-1β, IL-2, IL-3, IL-4, IL-5, IL-6, IL-9, IL-10, IL-12p40, IL-12p70, IL-13, IL-17, Eotaxin, G-CSF, GM-CSF, IFN-γ, KC, MCP-1, MIP-1α, MIP-1β, RANTES and TNF-α were included in this assay.

### Multiplex cytokine analysis on gestational tissue samples

#### Tissue collection

Uterus and placenta samples were collected at autopsy on gd17.5. A randomly selected longitudinal quarter of the uterus and a quarter of each placenta was snap frozen in liquid nitrogen for protein extraction.

#### Tissue preparation and cytokine analysis

Uterus and placenta tissue samples were processed using a Bio-Plex Cell Lysis Kit (BioRad Laboratories Inc), following the manufacturer instructions. Briefly, samples were homogenized with 500µl of lysing solution and frozen overnight at −80°C. The following day the samples were sonicated, centrifuged at 4500g for 4 mins and the supernatant collected. The protein content of the supernatant was determined using a *DC* Protein Assay (BioRad Laboratories Inc). 900µg/ml of protein was then assayed for cytokines using a separate Bio-Plex Pro mouse cytokine 23 plex kit (BioRad Laboratories Inc), following the manufacturer instructions.

### Tissue preparation and cytokine analysis

#### Tissue Collection

Uterus, decidua and placenta samples were collected at autopsy on gd17.5. The entire decidua and a randomly selected longitudinal quarter of the uterus was placed in RNAlater (Ambion) overnight at 4°C. Tissues were then transferred to a fresh tube and frozen at −80°C for RNA extraction and transcriptome profiling.

#### Tissue preparation, RNA extraction and transcriptome profiling

Decidua and uterus tissue samples were homogenized utilizing a rotor-stator homogenizer (Qiagen), and total RNA was extracted employing TRIzol (Ambion) followed by RNeasy MinElute (Qiagen). The integrity of the RNA was assessed on the Bioanalyzer (RIN: 9.4±0.2). Total RNA samples were shipped to AGRF for library preparation (TruSeq Stranded mRNA Library Prep Kit, Illumina) and sequencing (Illumina HiSeq2500, 50bp single-end reads, v4 chemistry, n=48). 25 million reads were generated per sample. The raw sequencing data is available from GEO (accession; XXYYZZ).

### RNA-Seq data analysis

The RNA-Seq data was analyzed in the R environment for statistical computing. The quality of the sequencing data was assessed with the Bioconductor package Rqc. Sequencing reads were aligned to the reference genome (mm10) employing Subread, and summarized at the gene level using featureCounts (57). Genes with less than 500 total counts across the data were removed from the analysis. Sample QC was performed by examining Relative Log Expression and Principal Component Analysis plots, and outlier samples were removed from the analysis. Differentially expressed genes (DEG) were identified employing negative binomial models in edgeR, with False Discovery Rate (FDR) control for multiple testing (58). The InnateDB database was utilized for pathways analysis (59). Molecular drivers of DEGs and networks were identified employing Upstream Regulator Analysis (30). Upstream regulators with absolute activation Z-scores < 2.0 were filtered out of the analysis, and were ranked by their overlap p-value. A coexpression network was constructed form the RNA-Seq data employing the WGCNA algorithm (28, 29). A separate network was constructed for each tissue (decidua, uterus). Prior to network analysis, the count data was transformed using the varianceStabilizingTransformation algorithm from the DESeq2 package (58). Network analysis was restricted to the top ~5,000 most variable genes, and these were identified using the varianceBasedfilter algorithm from the Bioconductor package DCGL. The modules identified by WGCNA were examined for enrichment of DEGs by calculating a median FDR for each module that was based on the gene-level statistics derived from the edgeR analysis.

### Statistical analysis

Statistical analyses were performed using GraphPad Prism software (version 6.0g for Mac OSX, La Jolia California USA). Unpaired t tests, one-way and two-way ANOVAs followed by Bonferroni multiple comparisons tests were used as indicated in the figure legends. The figures include the number of animals or litters per group.

Principle component analysis (PCA) was performed using the FactoMineR R package (19). Parameters used in the PCA included: clinical score, maternal weight, fetal weight, BAL cells and viral copy number (Supplementary Table 1). The first two dimensions (PCA 1 and PCA 2) accounted for 79% of the variability between samples.

## Author contributions

AB, PGH and DHS designed and supervised the study. NMS, JFLJ, KTM and MS performed the experiments. NMS, JFLJ, ACJ, KTM, NMT, JL, AB and DHS analyzed the data. SAR, SLP, CP contributed to project design, methodology and discussions on data interpretation. AB, PGH and DHS wrote the manuscript. All authors reviewed the final manuscript.

## Acknowledgments

We thank Prof. Casssandra Berry who kindly provided us with the Influenza virus. This study was funded principally by Nation Health and Medical Research Council (NHMRC) of Australia with supplementary support provided by OM Pharma (Geneva, Switzerland). AB is supported by a BrightSpark Foundation McCusker Fellowship (Western Australia, Australia). ACJ and KTM are recipients of an Australian Postgraduate Award and a Top-Up Award from the University of Western Australia.

## References

1. Lapinsky SE. Obstetric infections. Critical care clinics. 2013;29(3): 509-20.

2. Sappenfield E, Jamieson DJ, Kourtis AP. Pregnancy and susceptibility to infectious diseases. Infectious diseases in obstetrics and gynecology. 2013;2013:752852.

3. Kemp MW. Preterm birth, intrauterine infection, and fetal inflammation. Frontiers in immunology. 2014;5:574.

4. Romero R, Dey SK, Fisher SJ. Preterm labor: one syndrome, many causes. Science. 2014;345(6198): 760-5.

5. Arck PC, Hecher K. Fetomaternal immune cross-talk and its consequences for maternal and offspring’s health. Nature medicine. 2013;19(5): 548-56.

6. Barker DJ, Thornburg KL. The obstetric origins of health for a lifetime. Clinical obstetrics and gynecology. 2013;56(3): 511-9.

7. Meijer WJ, van Noortwijk AG, Bruinse HW, Wensing AM. Influenza virus infection in pregnancy: a review. Acta Obstet Gynecol Scand. 2015;94(8): 797-819.

8. Luan H, Zhang Q, Wang L, Wang C, Zhang M, Xu X, et al. OM85-BV induced the productions of IL-1beta, IL-6, and TNF-alpha via TLR4- and TLR2-mediated ERK1/2/NF-kappaB pathway in RAW264.7 cells. Journal of interferon & cytokine research: the official journal of the International Society for Interferon and Cytokine Research. 2014;34(7): 526-36.

9. Parola C, Salogni L, Vaira X, Scutera S, Somma P, Salvi V, et al. Selective activation of human dendritic cells by OM-85 through a NF-kB and MAPK dependent pathway. PLoS One. 2013;8(12):e82867.

10. Schaad UB. OM-85 BV, an immunostimulant in pediatric recurrent respiratory tract infections: a systematic review. World journal of pediatrics: WJP. 2010;6(1): 5-12.

11. Weinberger M. Can we prevent exacerbations of asthma caused by common cold viruses? The Journal of allergy and clinical immunology. 2010;126(4): 770-1.

12. Emmerich B, Emslander HP, Pachmann K, Hallek M, Milatovic D, Busch R. Local immunity in patients with chronic bronchitis and the effects of a bacterial extract, Broncho-Vaxom, on T lymphocytes, macrophages, gamma-interferon and secretory immunoglobulin A in bronchoalveolar lavage fluid and other variables. Respiration; international review of thoracic diseases. 1990;57(2): 90-9.

13. Collet JP, Ducruet T, Kramer MS, Haggerty J, Floret D, Chomel JJ, et al. Stimulation of nonspecific immunity to reduce the risk of recurrent infections in children attending day-care centers. The Epicreche Research Group. The Pediatric infectious disease journal. 1993;12(8): 648-52.

14. Razi CH, Harmanci K, Abaci A, Ozdemir O, Hizli S, Renda R, et al. The immunostimulant OM-85 BV prevents wheezing attacks in preschool children. The Journal of allergy and clinical immunology. 2010;126(4): 763-9.

15. Soler M, Mutterlein R, Cozma G. Double-blind study of OM-85 in patients with chronic bronchitis or mild chronic obstructive pulmonary disease. Respiration; international review of thoracic diseases. 2007;74(1): 26-32.

16. Strickland DH, Fear V, Shenton S, Wikstrom ME, Zosky G, Larcombe AN, et al. Persistent and compartmentalised disruption of dendritic cell subpopulations in the lung following influenza A virus infection. PLoS One. 2014;9(11):e111520.

17. Raj RS, Bonney EA, Phillippe M. Influenza, Immune System, and Pregnancy. Reproductive sciences. 2014.

18. Rasmussen SA, Jamieson DJ, Uyeki TM. Effects of influenza on pregnant women and infants. American journal of obstetrics and gynecology. 2012;207(3 Suppl):S3-8.

19. Lê S, Josse J, Husson F. FactoMineR: An R Package for Multivariate Analysis. 2008;25(1): 18.

20. Nadeau-Vallee M, Quiniou C, Palacios J, Hou X, Erfani A, Madaan A, et al. Novel Noncompetitive IL-1 Receptor-Biased Ligand Prevents Infection- and Inflammation-Induced Preterm Birth. Journal of immunology. 2015;195(7): 3402-15.

21. Robertson SA, Skinner RJ, Care AS. Essential role for IL-10 in resistance to lipopolysaccharide-induced preterm labor in mice. Journal of immunology. 2006;177(7): 4888-96.

22. Cappelletti M, Della Bella S, Ferrazzi E, Mavilio D, Divanovic S. Inflammation and preterm birth. Journal of leukocyte biology. 2016;99(1): 67-78.

23. Erlebacher A. Immunology of the Maternal-Fetal Interface. Annual Review of Immunology. 2013;31(1): 387-411.

24 Robertson SA, Petroff MG, Hunt JS. Immunology of Pregnancy. In: Plant TM, Zeleznik AJ, editors. Knobil and Neill’s Physiology of Reproduction. 4th ed. Oxford, UK: Elsevier; 2015. p. 1835-1874.

25. Tagliani E, Erlebacher A. Dendritic cell function at the maternal-fetal interface. Expert review of clinical immunology. 2011;7(5): 593-602.

26. Zhao H, Kalish F, Schulz S, Yang Y, Wong RJ, Stevenson DK. Unique roles of infiltrating myeloid cells in the murine uterus during early to midpregnancy. Journal of immunology. 2015;194(8): 3713-22.

27. Collins MK, Tay CS, Erlebacher A. Dendritic cell entrapment within the pregnant uterus inhibits immune surveillance of the maternal/fetal interface in mice. The Journal of clinical investigation. 2009;119(7): 2062-73.

28. Bosco A, McKenna KL, Firth MJ, Sly PD, Holt PG. A network modeling approach to analysis of the Th2 memory responses underlying human atopic disease. Journal of immunology. 2009;182(10): 6011-21.

29. Langfelder P, Horvath S. WGCNA: an R package for weighted correlation network analysis. BMC bioinformatics. 2008;9:559.

30. Krämer A, Green J, Pollard J, Jr. Tugendreich S. Causal analysis approaches in Ingenuity Pathway Analysis. Bioinformatics (Oxford, England). 2014;30(4): 523-30.

31. Pazos M, Sperling RS, Moran TM, Kraus TA. The influence of pregnancy on systemic immunity. Immunologic research. 2012;54(1–3):254-61.

32. Levy O, Wynn JL. A prime time for trained immunity: innate immune memory in newborns and infants. Neonatology. 2014;105(2): 136-41.

33. Meissner HC. Viral Bronchiolitis in Children. N Engl J Med. 2016;374(1): 62-72.

34. Hacker H, Redecke V, Blagoev B, Kratchmarova I, Hsu LC, Wang GG, et al. Specificity in Toll-like receptor signalling through distinct effector functions of TRAF3 and TRAF6. Nature. 2006;439(7073): 204-7.

35. Kagan JC, Su T, Horng T, Chow A, Akira S, Medzhitov R. TRAM couples endocytosis of Toll-like receptor 4 to the induction of interferon-beta. Nature immunology. 2008;9(4): 361-8.

36. Parker D, Prince A. Type I interferon response to extracellular bacteria in the airway epithelium. Trends in immunology. 2011;32(12): 582-8.

37. Blank V, Hirsch E, Challis JR, Romero R, Lye SJ. Cytokine signaling, inflammation, innate immunity and preterm labour - a workshop report. Placenta. 2008;>29 Suppl A:S102-4.

38. Challis JR, Lockwood CJ, Myatt L, Norman JE, Strauss JF, 3rd, Petraglia F. Inflammation and pregnancy. Reproductive sciences. 2009;16(2): 206-15.

39. Husebye H, Halaas O, Stenmark H, Tunheim G, Sandanger O, Bogen B, et al. Endocytic pathways regulate Toll-like receptor 4 signaling and link innate and adaptive immunity. EMBO J. 2006;25(4): 683-92.

40. van der Meer JW, Joosten LA, Riksen N, Netea MG. Trained immunity: A smart way to enhance innate immune defence. Molecular immunology. 2015;68(1): 40-4.

41. Kleinnijenhuis J, Quintin J, Preijers F, Joosten LA, Jacobs C, Xavier RJ, et al. BCG-induced trained immunity in NK cells: Role for non-specific protection to infection. Clinical immunology. 2014;155(2): 213-9.

42. von Mutius E, Radon, K.,. Living on a farm: impact on asthma induction and clinical course. Immunol Allergy Clin North Am. 2008;28(3): 631-47.

43. von Mutius E. The microbial environment and its influence on asthma prevention in early life. The Journal of allergy and clinical immunology. 2016;137(3): 680-9.

44. Jahnsen FL, Strickland DH, Thomas JA, Tobagus IT, Napoli S, Zosky GR, et al. Accelerated antigen sampling and transport by airway mucosal dendritic cells following inhalation of a bacterial stimulus. Journal of immunology. 2006;177(9): 5861-7.

45. Platt AM, Randolph GJ. Dendritic cell migration through the lymphatic vasculature to lymph nodes. Advances in immunology. 2013;120:51–68.

46. Asselin-Paturel C, Brizard G, Chemin K, Boonstra A, O’Garra A, Vicari A, et al. Type I interferon dependence of plasmacytoid dendritic cell activation and migration. The Journal of experimental medicine. 2005;201(7): 1157-67.

47. Dai J, Megjugorac NJ, Amrute SB, Fitzgerald-Bocarsly P. Regulation of IFN regulatory factor-7 and IFN-alpha production by enveloped virus and lipopolysaccharide in human plasmacytoid dendritic cells. Journal of immunology. 2004;173(3): 1535-48.

48. Amit I, Garber M, Chevrier N, Leite AP, Donner Y, Eisenhaure T, et al. Unbiased reconstruction of a mammalian transcriptional network mediating pathogen responses. Science. 2009;326(5950): 257-63.

49. Bogdan C, Mattner J, Schleicher U. The role of type I interferons in non-viral infections. Immunological reviews. 2004;202:33–48.

50. Pasquali C, Salami O, Taneja M, Gollwitzer ES, Trompette A, Pattaroni C, et al. Enhanced Mucosal Antibody Production and Protection against Respiratory Infections Following an Orally Administered Bacterial Extract. Frontiers in medicine. 2014;1:41.

51. Bizargity P, Bonney EA. Dendritic cells: a family portrait at mid-gestation. Immunology. 2009;126(4): 565-78.

52. Gomez-Lopez N, Olson DM, Robertson SA. Interleukin-6 controls uterine Th9 cells and CD8(+) T regulatory cells to accelerate parturition in mice. Immunology and cell biology. 2016;94(1): 79-89.

53. Merad M, Sathe P, Helft J, Miller J, Mortha A. The dendritic cell lineage: ontogeny and function of dendritic cells and their subsets in the steady state and the inflamed setting. Annu Rev Immunol. 2013;31:563–604.

54. Segura E, Amigorena S. Inflammatory dendritic cells in mice and humans. Trends in immunology. 2013;34(9): 440-5.

55. Diggins KE, Ferrell PB, Jr. Irish JM. Methods for discovery and characterization of cell subsets in high dimensional mass cytometry data. Methods (San Diego, Calif). 2015;82:55–63.

56. Kotecha N, Krutzik PO, Irish JM. Web-based analysis and publication of flow cytometry experiments. Current protocols in cytometry/editorial board, J Paul Robinson, managing editor [et al]. 2010;Chapter 10:Unit 10.7.

57. Liao Y, Smyth GK, Shi W. The Subread aligner: fast, accurate and scalable read mapping by seed-and-vote. Nucleic acids research. 2013;41(10):e108.

58. Anders S, McCarthy DJ, Chen Y, Okoniewski M, Smyth GK, Huber W, et al. Count-based differential expression analysis of RNA sequencing data using R and Bioconductor. Nature protocols. 2013;8(9): 1765-86.

59. Lynn DJ, Winsor GL, Chan C, Richard N, Laird MR, Barsky A, et al. InnateDB: facilitating systems-level analyses of the mammalian innate immune response. Molecular systems biology. 2008;4:218.

